# Ethambutol resistance in *Mycobacterium avium-intracellulare* is associated with mutations in the *embA–embB* promoter and *embB*

**DOI:** 10.64898/2026.05.23.727351

**Authors:** Asami Osugi, Keiji Fujiwara, Masashi Ito, Yu Kurahara, Kozo Morimoto, Satoshi Mitarai

**Author notes:** Address correspondence to Asami Osugi.

## Abstract

Ethambutol (EMB) is a vital drug for treating *Mycobacterium avium-intracellulare* (MAI) infections; however, the genomic mutations underlying EMB resistance in MAI remain unclear. Herein, we evaluated eight sets of MAI clinical isolates, each containing at least two serial isolates collected from the same patient who received EMB in Japan. In four sets, the isolates independently increased EMB MIC by 4-fold, coinciding with mutations in the upstream region of *embA* or those corresponding to *Mycobacterium tuberculosis* (Mtb) *embB* Met306Val and Gln497Arg. Based on the increased EMB MIC values, we defined normal and elevated EMB MICs as ≤8 µg/mL and ≥16 µg/mL, respectively. In the other four sets, all of the isolates had elevated EMB MICs. *In silico* promoter prediction and expression analysis indicated that the upstream region of *embA* corresponds to the *embA–embB* promoter region, and mutations in this region increased the transcription of *embA* and *embB*, increasing EMB MICs. Furthermore, the analysis of 60 epidemiologically unrelated strains revealed that isolates with mutations in the *embA–embB* promoter and at *embB* codons 306/497 exhibited significantly higher EMB MICs compared with those without mutations. Publicly available genomic data demonstrate the worldwide occurrence of these mutations in clinical isolates. These results establish an association between elevated EMB MICs and mutations at *embB* codons 306/497 and the *embA–embB* promoter and are expected to predict EMB resistance.

## Introduction

The incidence of lung disease caused by nontuberculous mycobacterial infection has been increasing globally over the past few decades, with *Mycobacterium avium-intracellulare* (MAI), a member of the *Mycobacterium avium-intracellulare* complex (MAC), being the most common causative species (1-3). Although macrolides are considered the most effective pharmacological treatments, MAI lung disease is challenging to treat and often results in failure (4). Since ethambutol (EMB) has been considered to repress the emergence of macrolide-resistant bacteria, a three-drug combination comprising a macrolide, rifampicin, and EMB has been recommended for treating macrolide-susceptible MAC (5).

The mode of action of EMB and the associated acquired resistance mechanisms are relatively well understood in *Mycobacterium tuberculosis* (Mtb). Specifically, the *emb* gene cluster comprising *embC, embA*, and *embB* represents the primary target of EMB (6). Its protein products, the EmbA/EmbB heterodimers and EmbC/EmbC homodimers, catalyze the synthesis of their mycobacteria-specific cell wall components, mycolyl-arabinogalactan-peptidoglycan and lipoarabionomannan, respectively (7,8). EMB inhibits cell wall assembly by binding directly to EmbB and EmbC (8). Accordingly, WHO recognizes 13 *embB* resistance mutations affecting the following six codons in a decreasing order of frequency: Met306Val/Ile/Leu, Gln497Arg/Lys, Gly406Ala/Asp/Ser/Cys, Asp354Ala, Tyr319Ser/Cys, and Asp328Tyr (9-11). Moreover, other rarer resistance mechanisms likely exist that are not currently endorsed by WHO. These include mutations upstream of *embA* and *embC* that result in the overexpression of the *embA–embB* operon and *embC*, respectively (9-13). Finally, *ubiA*, which synthesizes the precursors of reactions catalyzed by EmbA/EmbB and EmbC/EmbC, has also been implicated in EMB resistance (14,15). EMB-resistant Mtb strains may harbor multiple resistance mutations, given that most, if not all, of these mechanisms confer only modest MIC increases (13,14). Conversely, the effects of mutations in the *emb* gene cluster on EMB resistance in MAI have not been well studied to date.

Despite recommendations for clinical use, the Clinical and Laboratory Standards Institute (CLSI) did not establish a breakpoint for drug susceptibility testing (DST) of EMB for MAC, because previous studies showed no correlation between *in vitro* MICs and response in patients with MAC (13). One potential explanation for this lack of correlation is that EMB offers little or no clinical benefit (16). In fact, EMB MICs in MAC may be too high for EMB to be clinically effective at the current dose (17). Moreover, Mtb has a breakpoint established by the CLSI (15); however, it is not conclusive because no correlation with clinical outcome, as in MAI, has been established, and low reproducibility of DST has been observed (10,14). Therefore, DST of EMB may also be less reproducible in MAI.

This study attempts to characterize mutations that increase the EMB MIC in MAI. To the best of our knowledge, this work provides the first comprehensive identification of mutations associated with the EMB MIC in MAI.

## Results

### Analysis of longitudinal clinical isolates

To identify mutations associated with EMB MICs in MAI, we analyzed eight sets of longitudinal clinical isolates that received EMB treatment in Japan (Tables 1, S1, and S2). Of the eight patients analyzed, four had a 4-fold increase in MIC between isolates in at least 4 of 6 independent experiments. Whole-genome sequencing revealed that mutations in the *emb* gene cluster were associated with an increase in MIC in these isolates. In patients 3, 9, and 12, *embB* mutations corresponding to Mtb Met306Val and Gln497Arg, which are associated with EMB resistance in Mtb (9) or those upstream of *embA*, which were suggested for EMB resistance in Mtb (10-12,18,19), were detected in the isolates with the highest MIC value in each set, but not in those with the lowest MIC value. In patient 5, the isolate with the lowest MIC harbored *embB* Met306Val, and allele frequencies (AFs) with an additional mutation upstream of *embA* were increased from 0.12 to 1, which coincided with an increase in MIC, indicating that AF 0.12 was too low to increase the MIC in our conditions. Because any mutation in the four patients conferred an MIC of ≥16 µg/mL, an elevated MIC was defined as ≥16 µg/mL, and a normal MIC was designated ≤8 µg/mL EMB in the present study. In the other four patients, all isolates had MICs ≥ 16 µg/mL and harbored more than one of the following mutations: *embB* Met306Val, *embB* Gln497Arg, and mutations upstream of *embA*. Two mutations were detected in all isolates exhibiting an MIC > 16 µg/mL among all six independent experiments. We evaluated the mutations in the 100-bp upstream and coding regions of *embA, embB, embC, embR*, and *ubiA*, which are known EMB resistance genes in Mtb, but none correlated with the EMB MICs, except for *embB* Met306Val, *embB* Gln497Arg, and mutations upstream of *embA*.

**Table 1.** *embB* Met306Val, *embB* Gln497Arg, and *embA* upstream mutations correlated with elevated ethambutol MICs from MAI isolates longitudinally collected from patients with chronic infections. Isolates with MICs ≤8 µg/mL were considered to have normal MICs, whereas those with predominantly ≥16 µg/mL were classified to have elevated MICs. In the increased group, 4-fold increase in MICs coincided with mutations. In the elevated group, all isolates had elevated MICs, and no change was observed in MICs or mutations.

| Group | Patient | Sample | MIC ($\mu\text{g/mL}$)<sup>a</sup> | Mutation<sup>b</sup> | | | |
| --- | --- | --- | --- | --- | --- | --- | --- |
|  |  |  |  | *embB* coding<sup>c</sup> | AF | *embA* upstream | AF |
| Increase | 3 | 1 | 4 | Met306Val<sup>d</sup><br>(275) | N.D. | N.A. |  |
|  |  | 2 | 4 |  | N.D. |  |  |
|  |  | 3 | 8 |  | N.D. |  |  |
|  |  | 4 | 16 |  | 1 |  |  |
|  | 5 | 1 | 16 | Met306Val<sup>d</sup><br>(275) | 0.99 | G-11A | 0.12 |
|  |  | 2 | 32 |  | 1 |  | 1 |
|  |  | 3 | 32 |  | 1 |  | 1 |
|  |  | 4 | 32 |  | 1 |  | 1 |
|  |  | 5 | 64 |  | 1 |  | 1 |
|  | 9 | 1 | 4 | N.A. |  | C-12T | N.D |
|  |  | 2 | 4 |  |  |  | N.D |
|  |  | 3 | 16 |  |  |  | 1 |
|  |  | 4 | 16 |  |  |  | 0.99 |
|  | 12 | 1 | 4 | Gln497Arg<sup>d</sup><br>(466) | N.D. | C-16G | N.D. |
|  |  | 2 | 4 |  | N.D. |  | N.D. |
|  |  | 3 | 32 |  | 0.94 |  | 0.99 |
| Elevated | 11 | 1 | 32 | Met306Val<sup>d</sup><br>(275) | 0.99 | T-15G | 0.97 |
|  |  | 2 | 64 |  | 1 |  | 1 |
|  |  | 3 | 64 |  | 1 |  | 1 |
|  |  | 4 | 64 |  | 1 |  | 0.99 |
|  | 17 | 1 | 16 | Gln497Arg<sup>d</sup><br>(466) | 0.97 | N.A. |  |
|  |  | 2 | 16 |  | 0.99 |  |  |
|  | 29 | 1 | 16 | N.A. |  | G-11A | 1 |
|  |  | 2 | 16 |  |  |  | 1 |
|  | 30 | 1 | 16 | N.A. |  | C-12T | 1 |
|  |  | 2 | 16 |  |  |  | 1 |
|  |  | 3 | 16 |  |  |  | 1 |

<sup>a</sup> Values in the representative experiment from six independent experiments is shown.
<sup>b</sup> AF, allele frequency. N.A., not applicable. N.D., not detected using an AF of 0.05 as the cut-off.
<sup>c</sup> The first number corresponds to the amino acid position in Mtb, whereas the number in parentheses
indicates the position in MAI.
<sup>d</sup> EMB resistance mutation in the WHO mutation catalog for Mtb.

### Comparison of the *emb* gene cluster in MAI with Mtb

Because associations between mutations in the *emb* gene cluster and elevated MICs were observed in MAI, as in Mtb, the gene orders and sequences were compared between species (Figure 1). The analysis of the genomes of the MAI strains 104 and OCU901 yielded similar results. In the Mtb reference strain H37Rv genome, the *emb* gene cluster consists of *embC, embA*, and *embB* in this order, with its transcriptional regulator *embR* located at a distant site on the chromosome. Conversely, the MAI *emb* gene cluster consists of *embC, embR, embA*, and *embB* in order. In the Mtb, the upstream region of *embA* contains the promoter for *embA* and *embB*, which are co-transcribed (18), and 39/50 bps upstream of *embA* were conserved between Mtb and MAI, despite the insertion of *embR* in MAI. All four mutations observed in our longitudinal isolates occurred in the region corresponding to the −10 box, a component of the bacterial promoter. Furthermore, 7/9 bp inside the −10 box were conserved between the two species, and four mutations were identified in two conserved and two non-conserved positions.

**Figure 1.**
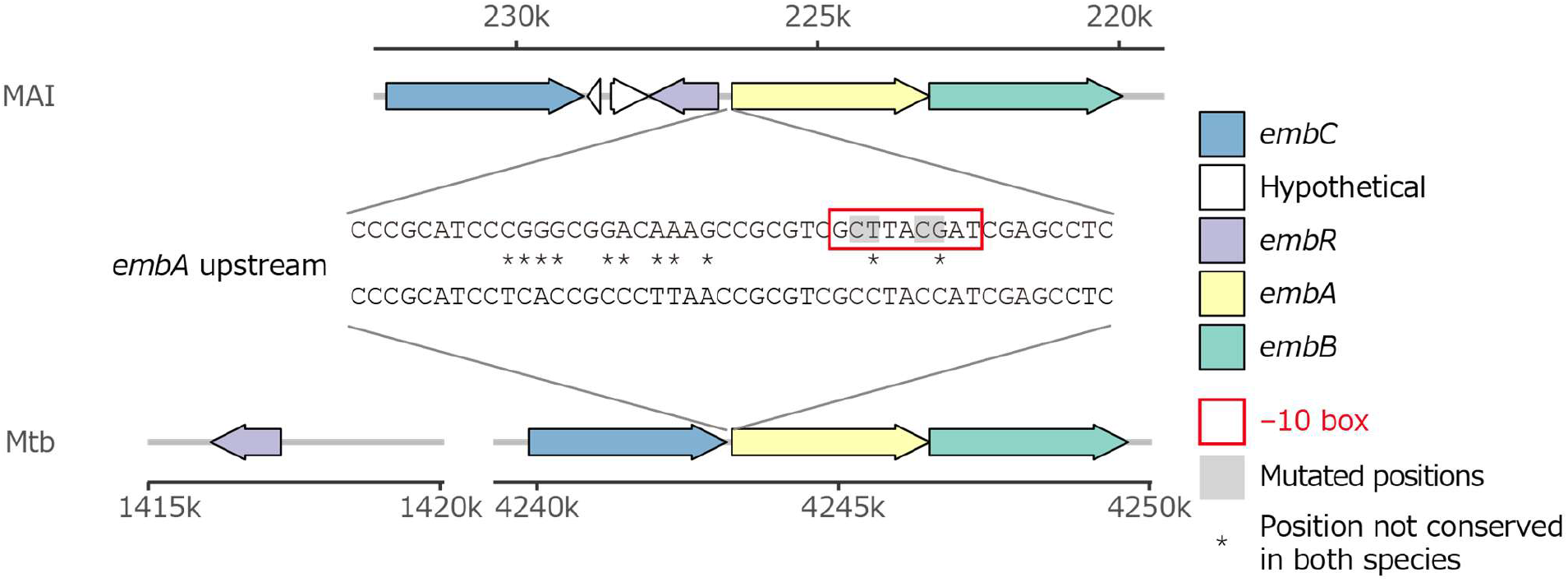
Co-key ethambutol resistance genes in MAI and Mtb. Genes are shown as arrows, with their coding orientation indicated by the arrow direction. The −10 box was predicted by the BPROM software in the presence of the G11A mutation (Table 2). Gray squares represent the mutated positions described in Table 1.

### Evaluation of the effect of mutations upstream of *embA*

Because mutations in the −10 box can affect the activity of a promoter, we determined the effect of the four mutations upstream of *embA* from the longitudinal isolates (Table 2). SAPPHIRE predicts the promoters in three versions: model-trained versions on dsRNA-seq data from *Pseudomonas aeruginosa* and *Salmonella enterica*, as well as an untrained version (20). All four mutations decreased the *p*-value for promoter prediction in both model-training versions, suggesting increased promoter activity *in silico*. SAPPHIRE without model training did not predict promoters, whereas BPROM predicted promoters only with the mutation at 11 bp upstream of *embA*. Therefore, we concluded that the upstream region of *embA* likely corresponds to the *embA–embB* promoter in MAI, and their mutations affected the promoter activities.

**Table 2.** Mutations upstream of *embA* in MAI increased the probabilities of promoter prediction *in silico*. Sequences in the *embA* upstream in MAI were analyzed using four *in silico* promoter prediction tools SAPPHIRE with three versions and BPROM. The values represent the *p*-values for the predictions if they were predicted. The Mtb sequence was also analyzed as a control. N.P. represents not predicted as a promoter.

| Species | *embA* promoter mutation<sup>a</sup> | *p*-value for the prediction in SAPPHERE |  |  | BPROM |
| --- | --- | --- | --- | --- | --- |
|  |  | CNN_salmonella | CNN_pseudomonas | No CNN |  |
| MAI | WT | 0.049 | 0.001 | N.P. | N.P. |
|  | G11A | 0.015 | 0.000 | N.P. | Predicted |
|  | C12T | 0.023 | 0.001 | N.P. | N.P. |
|  | T15G | 0.031 | 0.000 | N.P. | N.P. |
|  | C16G | 0.028 | 0.000 | N.P. | N.P. |
| Mtb | WT | 0.037 | 0.000 | N.P. | N.P. |

<sup>a</sup> WT is a sequence of the reference genome of the corresponding species. Other descriptions correspond to mutations shown in Table 1.

Next, we tested this hypothesis by measuring the expression of *embA* and *embB* in some of the longitudinal isolates (Figure 2). Since the basal expression between *M. avium* and *M. intracellulare* might differ, only *M. avium* samples were analyzed. The expression of both genes was significantly lower in isolates containing wild-type (WT) promoter sequences, including those with *embB* mutations (e.g., isolate 4 from patient 3 or isolate 4 from patient 12), compared with promoter mutants. Of the promoter mutants, the expression of both genes was significantly higher in mutants at positions that were not conserved with Mtb (i.e., G11A and T15G) compared with those that were conserved (i.e., C12T and C16G). Notably, we tested matched WT and mutant isolates from the same patients for three of the four mutations (i.e., G11A, C12T, and C16G), thereby increasing the likelihood that these mutations were causative, rather than merely associated with elevated expression.

**Figure 2.**
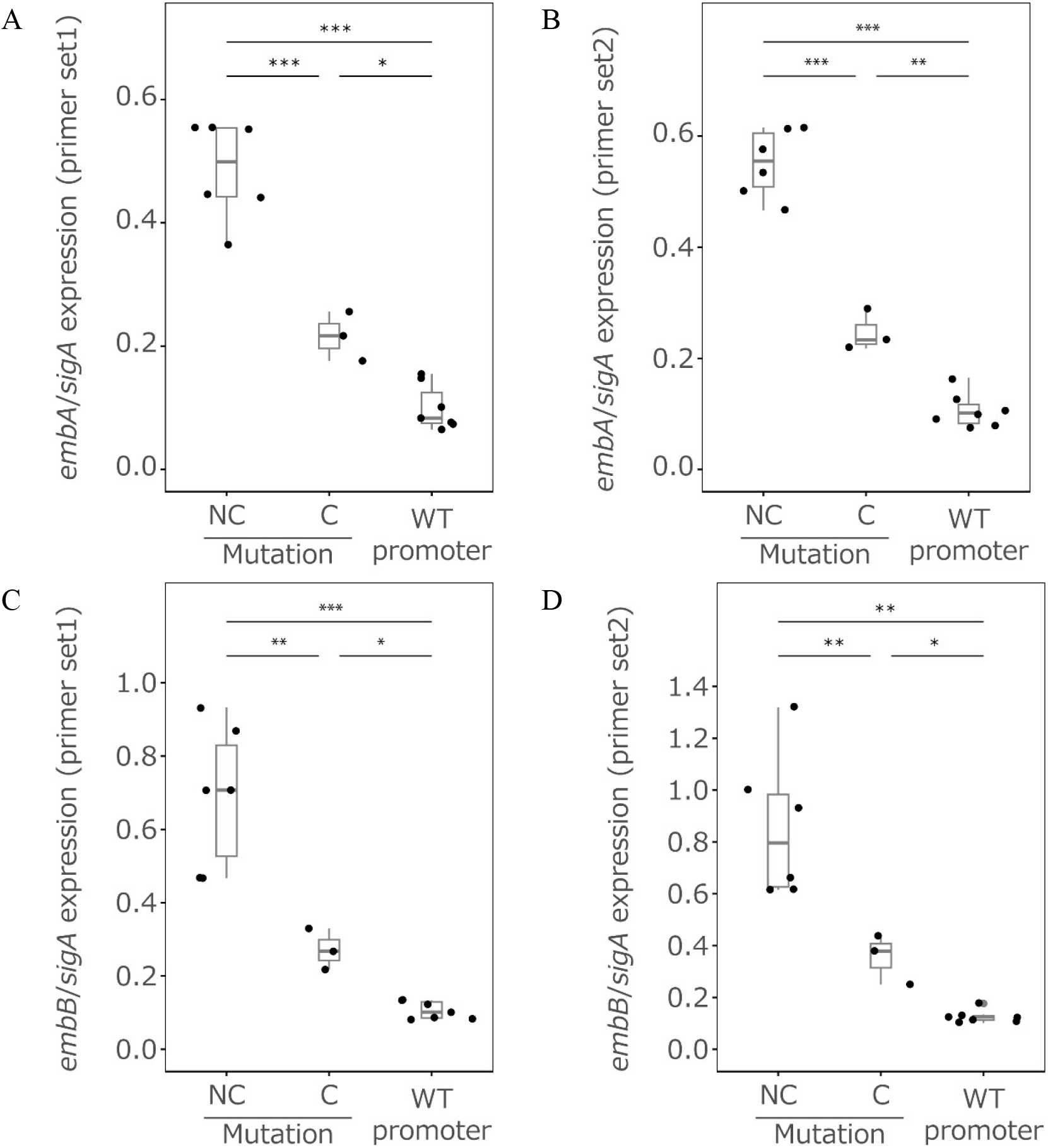
Promoter mutations correlated with increased expression of *embA* and *embB* in MAI. Quantitative real-time PCR was performed for (A and B) *embA* and (C and D) *embB* using two different primer sets for each gene. The longitudinally collected clinical isolates were divided into three groups based on their *embA* promoter sequence: WT and mutants with a change at a conserved (C) position between MAI and Mtb or non-conserved (NC) positions. The isolate list in each group is described in Table S1. \**p* < 0.05, \*\**p* < 0.01, and \*\*\**p* < 0.001 by *t*-test between the two groups.

### Analysis of 60 epidemiologically unrelated isolates

We evaluated correlations between EMB MICs and mutations in *embB* codons 306/497 and the *embA– embB* promoter in 60 epidemiologically unrelated isolates that were enriched for mutants identified during routine testing of clinical isolates with EMB MICs ≥16 µg/mL (Table S3). When MICs were measured using the same method as in Table 1, only 18/60 isolates had values ≥16 µg/mL, which was inconsistent with the routine testing (Figure 3, Table S3). An isolate harboring *embB* Met306Thr with an AF of 0.13 was considered to have no mutations, based on the results in which an AF of 0.12 was insufficient to elevate the MIC values (Table 1). Isolates with mutations in the *embA–embB* promoter and at *embB* codons 306/497 showed significantly higher MIC values compared with isolates without mutations, which confirms the correlation between the MICs and these mutations (Figure 3, Tables S3 and S6). Two isolates with an MIC ≤ 8 µg/mL had mutations in *embB* Gln497Lys or *embA* C16G. Except for the *embA–embB* promoter and *embB* codons 306/497, none of the mutations in the 100 bp upstream and coding regions of *embA, embB, embC, embR*, and *ubiA* were enriched in isolates with MICs ≥16 µg/mL (Table S4).

**Figure 3.**
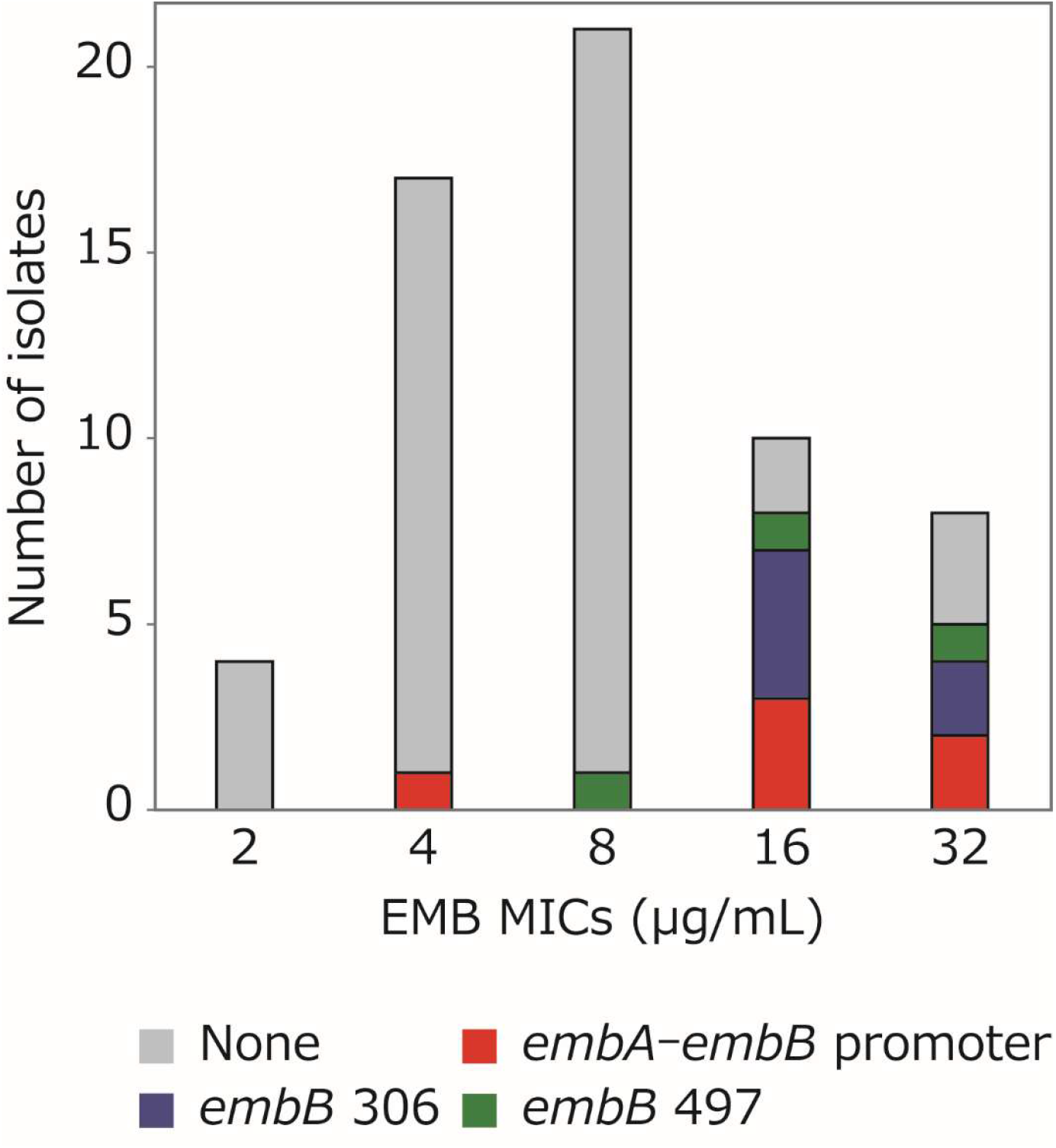
EMB MICs of 60 epidemiologically unrelated MAI isolates correlated with mutations in the *embA–embB* promoter and at *embB* codons 306/497. EMB MICs were significantly higher in isolates with mutations in the *embA–embB* promoter or at *embB* codons 306/497 (*p* < 0.0001, Mann–Whitney U test). Isolates were analyzed, and their MIC values are listed in Table S3.

### Analysis of public genomes from different hosts

Although the mutations we described above have not been well characterized in MAI, previously reported isolates may harbor these mutations. We analyzed 3,573 genomes from the Sequence Read Archive (SRA) (Tables S5 and S6). Mutations in the *embA–embB* promoter and at *embB* codons 306/497 were significantly enriched in isolates collected from humans compared with non-human sources, although drug treatment histories were not available (Table 3).

**Table 3.**
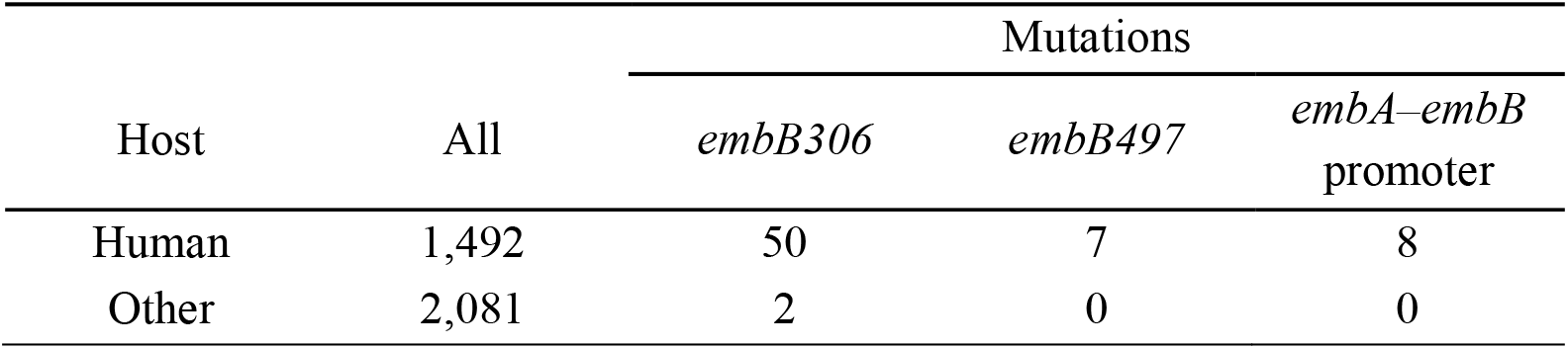
Mutations in the *embA–embB* promoter and *embB* codons 306/497 were overrepresented in MAI genomes isolated from humans. Publicly obtained genome data of 3,573 strains were analyzed. The mutations in the *embA–embB* promoter and at *embB* codons 306/497 were significantly enriched in humans than in others (*p* < 0.0001, Fisher’s exact test).

## Discussion

Despite the clinical significance of EMB, the molecular mechanisms underlying EMB resistance in MAI remain poorly understood. Herein, we found that mutations at embB codons 306/497 and in the *embA–embB* promoter region were associated with higher MICs in MAI (Tables 1 and 3). Of these mutations, those at *embB* codons 306/497 are considered resistance mutations in the Mtb WHO mutation catalog, but not those in the *embA–embB* promoter (9); however, involvement of the *embA– embB* promoter mutations in Mtb EMB resistance was suggested in several other studies (10-12,18, 19).

Based on EMB MIC distributions, most MAIs are resistant to therapeutic doses of EMB (17). Thus, the effects of EMB on MAI at therapeutic doses are unclear. In this study, four sets of MAI isolates evolved to exhibit a 4-fold increase in EMB MICs within the same patients who received EMB treatment (Table 1). These results suggest that elevating EMB MICs are beneficial to MAI, indicating the clinical efficacy of EMB therapy. Because EMB enhances the efficacy of other drugs, and combination therapy with macrolides and rifampicin is recommended for MAC (5,21), the therapeutic doses of EMB may be sufficient to enhance the efficacy of simultaneously administered drugs, although they are not sufficient for monotherapy, as indicated by the MIC values.

Among 42 epidemiologically unrelated isolates with EMB MICs ≤8 µg/mL, two had mutations in *embB* Gln497Lys or *embA* C-16G (Tables S3 and S4), which may be concordant with MICs ≤ 8 µg/mL. *embB* Gln497Lys was not observed in isolates with MICs ≥ 16 µg/mL in longitudinal and epidemiologically unrelated datasets, suggesting limited confidence in the resistance phenotype. *embA* C-16G was observed in an isolate with a MIC ≥ 16 µg/mL in the longitudinal dataset; however, its contribution to the MIC is unclear, because it occurred simultaneously with *embB* Gln497Arg. Other than these, no mutations in the *embA–embB* promoter or at *embB* codons 306/497 were detected in the 42 epidemiologically unrelated isolates with EMB MICs ≤ 8 µg/mL.

Previous studies in Mtb suggest that each mutation confers a slight increase in EMB MIC and combinations of mutations confer stronger resistance (11, 14). Our MAI isolates with multiple mutations also showed considerably high MICs. Isolates from patients 5 and 11 had EMB MICs of 64 µg/mL with mutations in the *embA–embB* promoter combined with *embB* Met306Val or Gln497Arg (Table 1). Additionally, a 4-fold MIC increase associated with each mutation was also observed in our study (Table 1). Such a moderate increases may result in technical variability of EMB MIC testing as reported in Mtb (13, 22). In fact, among 60 epidemiologically unrelated isolates with EMB MICs ≥ 16 µg/mL during routine testing, only 18 had MICs ≥ 16 µg/mL in three repeated tests, and the MIC values fluctuated slightly between independent experiments (Tables S1 and S3). Therefore, an alternative explanation for the lack of correlation between MICs and clinical outcome is that MIC testing cannot reliably distinguish between isolates containing normal and elevated MICs.

This study had two limitations. First, although the association between mutations in the *embA– embB* promoter, *embB* Met306, and *embB* Gln497 and increased MICs is plausible and supported by longitudinal and epidemiological data, we did not include genetically engineered transformants to directly prove causality. Second, none of the mutations in the *embA–embB* promoter and *embB* codons 306/497 were detected in five MAI isolates with an EMB MIC ≥ 16 µg/mL, indicating the presence of unknown mutations (Figure 3). We did not observe any relevant *ubiA* mutations that have recently been proposed as the dominant resistance mechanism in MAI (15). Therefore, further studies are needed to describe EMB-resistant mutations.

In the present study, mutations in the *embA–embB* promoter and at *embB* codons 306/497 coincided with EMB MICs ≥ 16 µg/mL. These results are expected to predict EMB MICs and subsequently, EMB efficacy in MAI.

## Materials and Methods

### Bacterial isolates

A total of 27 longitudinal MAI isolates from 8 patients and 60 epidemiologically unrelated MAI isolates from different patients were collected at the Fukujuji Hospital in Japan (Tables S1 and S3). Primary isolation was performed using 2% Ogawa medium (Kyokuto Pharmaceutical Industrial, Tokyo, Japan) or the automated BACTEC MGIT 960 system (Becton Dickinson, New Jersey, USA). The 60 epidemiologically unrelated MAI isolates were selected because each had an EMB MIC ≥ 16 µg/mL during routine testing using the BrothMIC SGM plate (Kyokuto Pharmaceutical Industrial), which is approved for diagnostic use in Japan by the Ministry of Health, Labor, and Welfare. Isolates were stored at −80°C and used for subsequent assays.

### DNA extraction and whole-genome sequencing

DNA was extracted as described previously (23). Libraries were prepared using the QIAseq FXDNA library kit (Qiagen, Venlo, Netherlands), and 150-bp paired-end sequencing was performed on the NextSeq 550 System (Illumina, California, USA).

### Analysis of genomic data

The sequence reads were aligned to the OCU901 genome (RefSeq genome assembly GCF_002716925.3) using the Burrows–Wheeler Aligner mem v0.7.17 (24). This reference was chosen because a greater proportion of reads aligned against it compared with other complete genomes, such as JCM30622, OCU873, ATCC13950, TH135, HP17, and 104 (RefSeq genome assemblies GCF_010731935.1, GCF_002716965.3, GCF_000277125.1, GCF_000829075.1, GCF_002716905.3, and GCF_000014985.1, respectively). Single-nucleotide polymorphisms were identified for each patient using bcftools mpileup and filtered for a mapping quality > 30, depth of high-quality bases > 20 in the FORMAT column, and QUAL column > 25 (25). The mutations in the coding and the 100-bp upstream regions of *embA, embB, embC, embR*, and *ubiA* (Genebank accession numbers: BJP78_RS01015, BJP78_RS01075, BJP78_RS01095, BJP78_RS01070, BJP78_RS01080, and BJP78_RS01010) were detected if AFs were >0.05.

### Promoter predictions

The 100-bp sequences upstream of *embA* were extracted from MAI strains 104 and OCU901 (RefSeq genome assemblies GCF_002716925.3 and GCF_000014985.1) and Mtb strain H37Rv (RefSeq genome assembly GCF_000195955.3), and the same sequences between the two MAI strains were confirmed. Four mutated sequences were generated *in silico* for MAI based on the mutations detected in the longitudinal isolates. In total, six sequences were applied for the promoter prediction using tools BPROM and three versions of SAPPHIRE (20, 26).

### Analysis of publicly available genomic data

Whole-genome sequencing data with *M. avium* complex (taxid 120793) were obtained from SRA (n = 4,783). The BioSample data containing a ScientificName of *Mycobacterium intracellulare subsp. chimaera, Mycobacterium colombiense, Mycobacterium marseillense, Mycobacterium timonense, Mycobacterium intracellulare subsp. yongonense, Mycobacterium arosiense, Mycobacterium mantenii, Mycobacterium bouchedurhonense*, or *Mycobacterium senriense*, were removed (n = 3,683; Table S5). All sequence data with one corresponding BioSample data, paired-end reads, aligned BAM file size > 100,000 B, and percentage aligned to OCU901 >55% were included in the analysis (n = 3,573; the filter column is NA in Table S5). Clinical isolates were identified based on the “host” information in the BioSample.

### MIC testing

Stored isolates were inoculated into Middlebrook 7H10 supplemented with 10% Middlebrook OADC (oleic acid-albumin-dextrose-catalase; Becton Dickinson), 0.5% (v/v) glycerol (Fujifilm Wako Pure Chem, Osaka, Japan). The cultured bacteria were subjected to MIC testing using the broth microdilution method. The MIC plate for EMB was prepared in Mueller–Hinton broth supplemented with 5% OADC using EMB dihydrochloride (MilliporeSigma, Massachusetts, USA), with EMB concentrations ranging from 0.5 to 512 µg/mL in 2-fold increments. MIC values were determined on the 7th day at the concentration at which bacterial growth was decreased visually. The MIC of ATCC 700898 determined on each testing day served as the control, whereas the MIC distribution for ATCC 700898 was 4–8 µg/mL, which was consistent with the data from a large multicenter study using a Sensititre SLOMYCOI broth microdilution plate (Thermo Fisher Scientific, Massachusetts, USA; 23).

### RNA extraction and expression analysis

Bacteria were cultured in Middlebrook 7H9 supplemented with 10% Middlebrook ADC (albumin– dextrose–catalase; Becton Dickinson), 0.2% (v/v) glycerol, and 0.05% (v/v) Tween 80 (Sigma-Aldrich, Missouri, USA). Centrifuged pellets were suspended in DNA/RNA Shield (Zymo Research, California, USA) when the optical density at 530 nm (OD_530_) ranged from 0.2 to 0.6. After bead-beating, total RNA was extracted with TRIzol (Thermo Fisher Scientific), according to the manufacturer’s protocol. Following treatment with TURBO DNase (Thermo Fisher Scientific), total RNA was purified using a Monarch Spin RNA Cleanup Kit (50 μg) (New England Biolabs, Massachusetts, USA). An equivalent of 500 ng total RNA was subjected to cDNA synthesis using a ReverTra Ace qPCR RT Master Mix with gDNA Remover (TOYOBO, Fukui, Japan). qPCR was performed using the primer sets listed in Table S7 with KOD SYBR qPCR Mix (TOYOBO). The relative expression value of the target gene to *sigA* was calculated using the 2Δ^Ct target gene−ΔCt *sigA*^ method.

## Acknowledgments

This work was supported by Janssen Pharmaceutical K.K. (grant number TMC207-NTM-GEN-01) and Japan Agency for Medical Research and Development (grant number JP23gm1610013). The funders had no role in study design, data collection, interpretation, or the decision to submit the work for publication. We thank Akio Aono for providing isolates from Fukujuji Hospital and Makiko Hosoya and Yoshiko Shimomura for supporting whole-genome sequencing. We appreciate Claudio U. Köser for supporting the editing of the manuscript.

## Contributions

A.O. designed and performed all experiments in the laboratories. A.O. designed and performed whole-genome data and statistical analyses. K.F., Y.I., Y.K., and K.M. collected clinical samples. K.F. performed prescreening of EMB resistance. A.O. visualized results and wrote the original draft. All authors reviewed and edited the manuscript. K.M. and S.M. supervised this study. S.M. acquired funding.

## Data Availability

Sequence data supporting the findings of this study have been deposited in the National Center for Biotechnology Information (NCBI) SRA database with accession numbers PRJNA1469383, PRJNA1469271, and PRJNA1189166.

## Supplemental Materials

Table S1. Longitudinal MAI isolates investigated in this study.

MAI isolates longitudinally collected from patients who received ethambutol are listed. Ethambutol MIC values of six independent experiments are shown, with an asterisk indicating representative data shown in Table 1. The group names are shown in the expression analysis group column for the isolates analyzed in Figure 2.

Table S2. Mutations detected in candidate genes involved in ethambutol resistance from longitudinal MAI isolates.

Mutations in the 100-bp upstream and coding region of *embA, embB, embC, embR*, and *ubiA* are listed if the mutations were detected in at least one isolate described in Table S1. The values in the isolate name column represent the allele frequencies, with blank cells indicating that the mutation was not detected in all isolates from the patient. The catalog_final and catalog_data columns represent values in the final confidence grading and variant columns of the *M. tuberculosis* mutation catalog (9), if the corresponding mutation is listed and the values in the final confidence grading column were ≥3 in the catalog.

Table S3. Epidemiologically unrelated MAI isolates examined in this study.

Epidemiologically unrelated 60 MAI isolates examined are listed. Ethambutol MIC values of three independent experiments are shown, with an asterisk indicating representative data shown in Table 3.

Table S4. Mutations detected in candidate genes involved in ethambutol resistance from 60 epidemiologically unrelated MAI isolates.

Mutations in the 100-bp upstream and coding region of *embA, embB, embC, embR*, and *ubiA* were listed if the mutations were detected in at least one isolate described in Table S3 The values in the isolate name column represent allele frequencies, whereas blank cells indicate that the mutation was not detected. The catalog_final and catalog_data columns represent values in the final confidence grading and variant columns in the *M. tuberculosis* mutation catalog (9), if the corresponding mutation is listed and the values in the final confidence grading column were ≥3 in the catalog.

Table S5. Publicly available 3,573 MAI genomic data examined in this study.

The 3,683 genomic data obtained from the SRA of the NCBI were listed, with the information in columns SRA, RUN, Location, and Hosts. After quality checks, 3,573 data described as NA in the filter column were processed to detect the mutations.

Table S6. Mutations detected in candidate genes involved in ethambutol resistance from the publicly available 3,573 MAI genomic data.

Mutations in the 100-bp upstream and coding region of *embA, embB, embC, embR*, and *ubiA* were listed if the mutations were detected in at least one isolate described in Table S5. The values in the isolate name column represent allele frequencies, whereas blank cells indicate that the mutation was not detected. The catalog_final and catalog_data columns represent values in the final confidence grading and variant columns in the *M. tuberculosis* mutation catalog (9), respectively, if the corresponding mutation is listed and the values in the final confidence grading column were ≥3 in the catalog.

Table S7. Primer sets used in reverse-transcription quantitative

PCR Primer sets used for the experiment shown in Table 2.

